# Viral replication and interferon responses in bronchial epithelia is enhanced by Th17 cells

**DOI:** 10.1101/2025.10.10.681711

**Authors:** Weston T. Powell, Tomasz J. Janczyk, Patricia C. dela Cruz, Basilin Benson, Elizabeth R. Vanderwall, Camille R. Gates, Lucille M. Rich, Maria P. White, Nyssa B. Samanas, Kourtnie Whitfield, Gail H. Deutsch, Teal S. Hallstrand, Matthew C. Altman, Jason S. Debley

## Abstract

**Rationale:** The impact of Th17 lymphocytes on epithelial responses to rhinovirus infection in asthma is poorly characterized.

**Methods:** Bronchial epithelial cells (BECs) from children with asthma were differentiated to an organotypic epithelium and primed via co-culture with healthy donor Th17 lymphocytes for 4 days prior to apical infection with human rhinovirus-16 (RV-16). RNA sequencing with WGCNA analysis was performed to identify modules of gene expression altered by Th17 priming or RV-16 infection in BECs or Th17 cells. Gene expression was correlated with viral copy number and with secreted protein levels.

**Results:** Analysis identified 4,030 genes grouped into 9 named modules with differential gene expression in BECs due to Th17 priming and viral infection. Modules with increased expression with Th17 priming and RV-16 infection included Interferon, MAP-kinase and TNF Signaling modules, while expression of Cilia structure/function and Metabolism modules were decreased. Th17 cells co-cultured with RV-16 infected BECs exhibited increased expression of an Interferon and Viral Response Module without detectable direct viral infection of Th17 cells.Increased expression of the Interferon Signaling in BECs and Interferon Response in Th17 cells was correlated with increased viral copy number in BECs. Th17 priming of BECs led to increased secretion of IFN-α, IFN-γ, and IL-1β following RV-16 as compared to BECs alone.

**Conclusions:** Th17 lymphocytes enhance epithelial interferon responses to RV-16 infection in bronchial epithelium from asthmatic children.

## Introduction

Infections with human rhinoviruses are the most common trigger of asthma exacerbations in children and adults.^2^ Asthma endotypes exist on a spectrum of overlapping features from type-2 (T2) to non-T2 endotypes.^3^ While the role of allergic airway inflammation mediating T2 asthma has been studied in greater detail, the mechanisms of airway inflammation and signaling in non-T2 endotypes is less well understood, particularly in the setting of viral infections. Non-T2 asthma is hypothesized to be in part driven by an IL-17-mediated pathway through Th17 lymphocytes leading to neutrophilic inflammation.^4^ While Th17 lymphocytes are known to arise in response to bacterial and fungal antigens, they also function in autoimmune and inflammatory conditions and coordinate neutrophilic inflammation during human rhinovirus infections.^5^ Co-culture of Th17 lymphocytes with bronchial fibroblasts leads to an increase in IL-17 production, as well as IL-6, IL-1β, and IL-23, however, the impact on gene expression by the epithelium was not investigated.^6^ Furthermore, the impact of Th17 stimulation of the epithelium on subsequent neutrophil function was not assayed. Similarly, while basolateral IL-17C stimulation increased CXCL1 production and IL-17C production following rhinovirus infection of bronchial epithelial cells (BECs), the impact on subsequent neutrophil activation or stimulation of the epithelium by Th17 cells prior to infection was not investigated.^7^ Increased Th17 cells and Th17-stimulated cytokines have been observed in the bronchial mucosa of adults with severe asthma with frequent exacerbations in association with increased epithelial neutrophils.^8^ Accordingly, airway epithelial-Th17 cell crosstalk may underpin a pathogenic mechanism in non-T2 asthma, particularly in the setting of viral-triggered exacerbations. However, how Th17 cell crosstalk with the airway epithelium alters innate immune and inflammatory responses to rhinovirus infection remains uninvestigated. As viral infections trigger exacerbations in both T2 and non-T2 clinically characterized asthma, the role of non-allergic inflammatory cells such Th17 cells in modulating epithelial responses to viral infections needs to be further studied to facilitate development of novel therapies. We hypothesized that Th17 cells in co-culture with human BECs from children with asthma would alter innate immune responses by both Th17 and BECs following rhinovirus infection.

## Methods

### BEC cultures and differentiation

BECs were obtained from children with asthma undergoing elective surgery under studies #12490 and #1596 approved by the Seattle Children’s Hospital Institutional Review Board as we have previously described.^9^ Parents of subjects provided written consent and children over 7 years of age provided assent. Basal cells were selected and proliferated under submerged culture conditions using PneumaCult™EX-Plus medium (Stemcell™). Passage 2-3 BECs were differentiated for≥21 days at an air-liquid interface (ALI) to generate organotypic, pseudostratified epithelial cultures.^9^*BEC-Immune Cell Co-culture*: Primary human Th17 cells (STEMCELL™) from a healthy donor were suspended (25,000 cells/mL) in a 50:50 mixture of PneumaCult™ ALI medium and RPMI 1640 with 10% FBS and added to the basolateral chamber of transwell cultures below differentiated BECs with media changed every 48 hours.

### RV-16 infection and sample collection

BECs were infected with human rhinovirus A16 (RV-A16, Source: ATCC®)) at a multiplicity of infection (MOI) of 0.5 on the apical surface for 2 hours at34°C as we have previously described.^10^ For BEC-Immune cell co-culture, infection on the apical surface occurred on day 4 after the start of Th17-BEC co-culture. (**Figure 1**) At 24 hours following RV-16 infection (or 5 days after co-culture initiation for uninfected cultures), Th17 cells were removed for RNA isolation and human primary neutrophils from a healthy donor (iQ Biosciences®; 25,000 cells/mL) in a 50:50 mixture of PneumaCult™-ALI medium and RPMI 1640 with 10% FBS were added to the basolateral chamber below differentiated BECs. 24 hours later neutrophils and BECs were harvested independently for RNA isolation, and media was collected.

**Figure 1.**
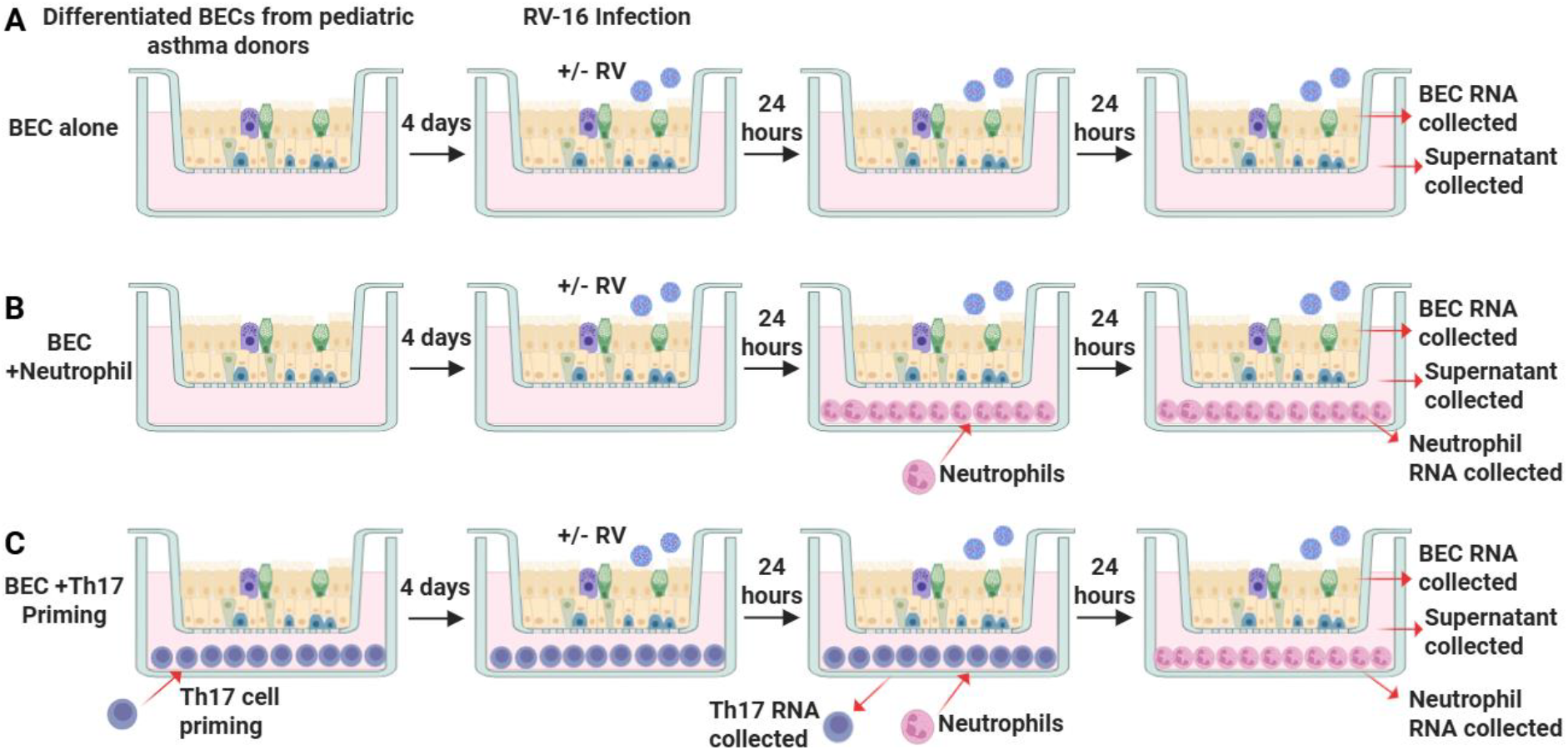
Schematic of experimental design for BEC alone cultures. **A.** BECs co-cultured with neutrophils for 24-hours. **B**. Th17 priming for BEC cells prior to neutrophil co-culture. Figure made with Biorender.

### Protein measurement

In collected media, protein concentrations of IL-17C, IL-1β, TNF-α, IFN-α, IFN-γ, CCL4, and CCL28 were measured via Human Luminex® Assay (R&D®) and CXCL10 and CXCL8 were measured using ELISA (Duoset R&D® DY266 and DY208).

### RNA sequencing

RNA was sequenced on a NextSeq 2000 sequencer (Illumina) with paired-end 53-base reads at a target depth of 5 million reads per sample, as previously described.^9^ Gene counts were generated as previously described using voomWithQualityWeights from the limma R package after alignment and quality-control.^9^ Samples that had human aligned counts greater than 1 million mapped reads and a median coefficient of variation coverage less than0.7 were kept for downstream analyses. Data from each cell type were processed separately. All raw bulk RNA-sequencing data will be uploaded to the NCBI Gene Expression Omnibus database at time of publication.

### Rhinovirus viral load quantification

To quantify rhinovirus viral load, the raw sequencing reads were aligned to the human reference genome (GRCh38) using STAR (v2.7.0a).^11^ Reads that were unmapped to the human reference genome (GRCh38) were used for viral quantification using the Salmon (v1.10.2) quasi-mapping algorithm against a custom viral reference database of 42 complete viral genomes (Supplemental Table 1).^12^ Transcript-level abundance estimates of number of reads were used to quantify rhinovirus abundance in each sample.

### Scanning electron microscopy

Transwells were fixed in 2% paraformaldehyde and 2% glutaraldehyde in 0.1M phosphate buffer at 4C°, then rinsed with 0.1M phosphate and post-fixed in 1% osmium tetroxide overnight at 4C° before being washed in water, dehydrated to 100% ethanol and critical point dried. Samples were then gold sputter-coated and imaged using a Thermo Fisher Scientific FEI Apreo VolumeScope scanning electron microscope.

### Statistics

Differentially expressed genes were identified using mixed effect linear models comparing expression differences by cell culture condition and infection status using the R package “kimma”.^13^ The model syntax was: gene_expression ∽ condition*infection and including a random effect for epithelial cell donorID. This identified 4,030 genes that reached a false discovery rate adjusted p-value (FDR) < 0.05 by the Benjamini-Hochberg correction procedure for cell culture condition. These genes were then utilized for supervised weighted gene co-expression network analysis (WGCNA) to identify genes with similar expression patterns and group them into modules. Module values were summarized by taking the mean of all genes (log2 transformed values) in the respective module. These modules were then modeled using the same linear model syntax to identify those showing differential expression patterns by cellculture condition and infection status at a stringent FDR < 0.05 for each pairwise comparison. Multiple hypothesis testing correction was performed using the Benjamini-Hochberg procedure. Pathway enrichment was performed using the clusterprofiler R package, which calculates a hypergeometric FDR corrected p-value for enrichment of public genesets.^14^ We used the gene ontology biological processes (GO_BP), KEGG, Reactome, Biocarta, and MSigDB Hallmark genesets for enrichment.^15–20^ The modules presented in the main text were annotated based on manual inspection of the enrichment terms and module genes. A similar model as used for gene expression was used for protein levels with multiple hypothesis testing correction using the Benjamini-Hochberg correction procedure. Pearson correlation of module expression in BECs to module expression in Th17 cells and for protein level to module expression was performed.

## Results

### BEC transcriptional responses to co-culture with Th17 cells demonstrate synergistic enhancement of a Type-II interferon signaling module

Primary BECs from 10 school age children with asthma (Table 1) were used in co-culture experiments. Cellular morphology, presence of cilia, and production of mucus were assessed visually under light microscopy of live cell cultures prior to initiation of co-culture experiments. Differentiation was also confirmed with formalin fixed paraffin embedded 5 micron hematoxylin and eosin stained sections and scanning electron microscopy. (**Figure S1**) We first investigated the impact of co-culture with neutrophils or Th17 lymphocytes then neutrophils on BEC transcriptional profiles with and without RV-16 infection. (**Figure 1**) Differential gene expression analysis identified 4,030 genes differentially expressed under co-culture conditions and 8,747 genes differentially expressed due to viral infection. These genes grouped into distinct co-expression modules that were differentially expressed among conditions and these modules were described based on gene set enrichment of the constituent genes. (**Figure 2**) Five named modules demonstrated an additive impact with significant differential expression due to Th17co-culture and RV infection. (**Figure 2A**) Three named modules showed an impact of co-culture with either neutrophils alone or with Th17 priming. (**Figure 2B**) One module had a synergistic effect of Th17 co-culture and viral infection with mean expression unchanged with co-culture alone, but enhanced expression after viral infection of Th17 co-cultured epithelium. (**Figure 2C**)

**Table 1.** Demographics of BEC donors for co-culture experiments.

|  |  |
| --- | --- |
| Sex (% Female) | 30% |
| Age (mean, range) | 11.2 (6.32-17.85) |
| FeNO (mean, range) | 20.3 (6-54) |
| FEV1 % (mean, range) | 102.1 (86-117) |
| FEV1/FVC (mean, range) | 0.82 (0.75-0.92) |
| IgE Level (mean, range) | 266.8 (3-1204) |
| % with history of severe exacerbation (ER visit, hospitalization or steroid use) | 90% |
| % with T2-high (FeNO >35ppb in steroid-naive or >20ppb on inhaled corticosteroid, or absolute eosinophils >300) | 40% |

**Figure 2.**
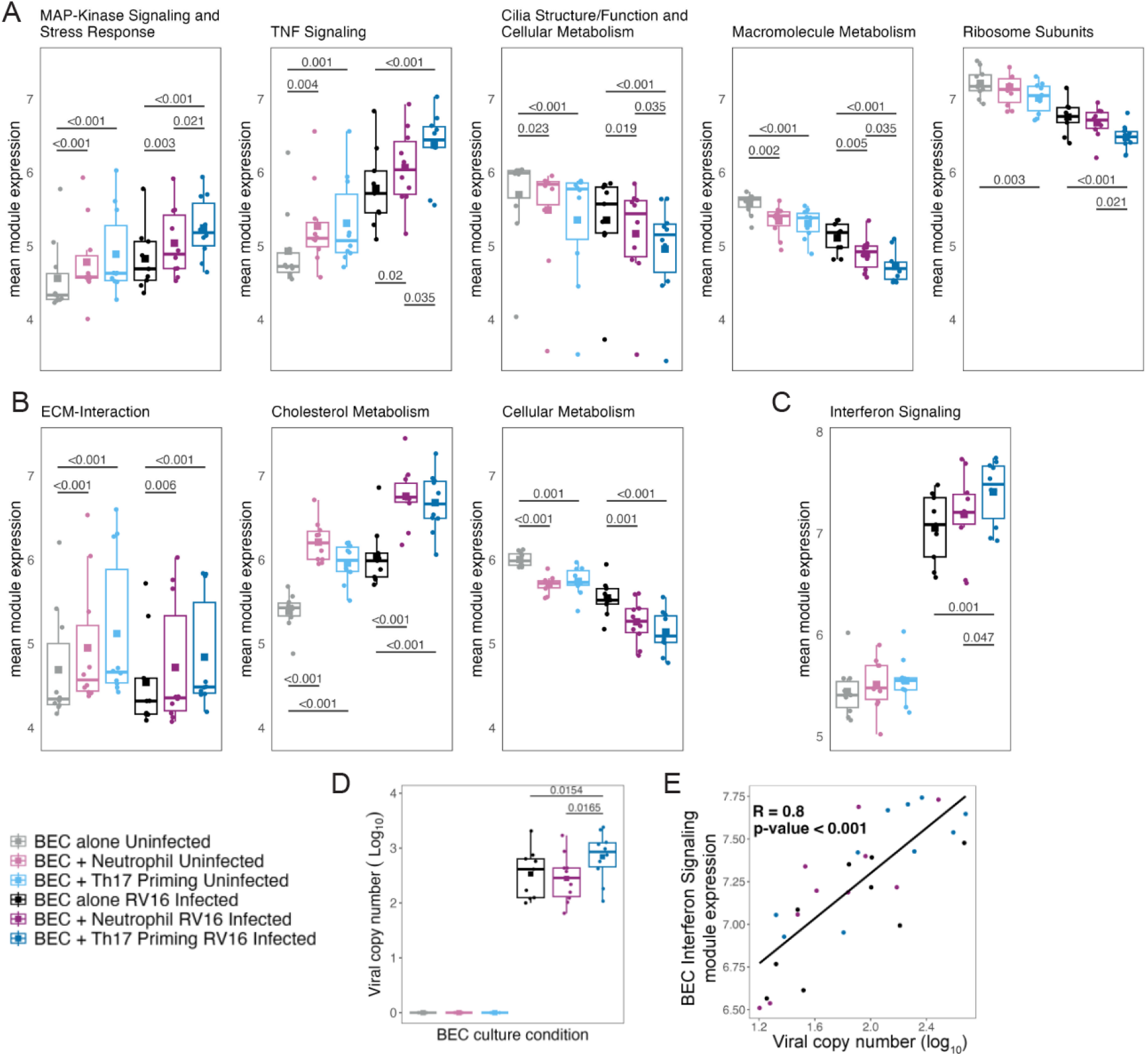
Th17 priming alters gene expression in BECs and viral copy number. **A.** 5 modules identified in BECs with additive differential gene expression from Th17 co-culture and RV-16 infection. **B**. 3 modules identified in BECs with differential expression from co-culture with neutrophils or Th17 cells. **C**. One module identified in BECs with mean gene expression unchanged due to co-culture under uninfected conditions, but synergistic increase in expression following infection of BECs primed with Th17 co-culture. **D**. Viral copy number in uninfected and infected BECs. **E**. Correlation of viral copy number with expression of the Interferon Signaling module in BECs across all infected culture conditions. All modules showed statistically significant differential expression when comparing uninfected to RV-16 infected conditions. For figure legibility, only the statistical comparisons within uninfected or within RV-16 infected conditions are shown.

Three modules showed decreased expression due to both Th17 co-culture and viral infection. (**Figure 2A**) The first module we termed “Cilia Structure/Function and Cellular metabolism” and was composed of 1531 genes which were predominantly cilia assembly/function genes, such as DYNLRB2 and *CCDC40*, and cellular metabolism genes such as *SOD1* and *NFIL3*. The second we termed “Macromolecule Metabolism” and was composed of 586 genes, such as *PTEN, POLR2B, EIF4A2, DNAJC8*, and *PRDX3*. The third gene module with decreased expression is termed “Ribosome Subunits and Viral Protein Translation” and was composed of 71 genes primarily related to ribosome function and protein translation.

Two modules exhibited increased expression in response to both Th17 co-culture and RV infection. We named one module “MAP-Kinase Signaling and Stress Response” which was composed of 323 genes, including *IL1A, IL1B, SERPINE1, CSF1, CCL4L2, IL17RA, TNF*, and *IL15*. The second module with increased expression was composed of 228 genes and named“TNF-signaling”, including genes such as *CSF3, MAP2K3, TRAF2, NFKB2, CXCL2, CXCL3, CXCL6, CXCL8, CCL20, CXCL6, IL18R1, IL6*, and *IRAK1*. (**Figure 2A**)

An “ECM-interaction” module and a “Cholesterol Metabolism” module exhibited increased expression in response to both RV infection and neutrophil co-culture, whereas a “Cellular Metabolism” module demonstrated decreased expression with both RV infection and neutrophil co-culture. (**Figure 2B**)

One module that showed synergistic increased expression from Th17 co-culture and RV-16 infection was composed of 45 genes, primarily Type II interferon signaling and viral response genes including *SP100, SERPINB1, STAT1, TRIM31, CXCL11, IFI44, GBP1, ISG20* and*HERC5*, and was thus named “Interferon Signaling”. (**Figure 2C)**

### Viral copy number mediation of BEC gene expression with Th17 priming

Viral load was increased in BECs primed with Th17 lymphocytes as compared to primary BECs alone or BECs co-cultured with neutrophils. (**Figure 2D**) Viral genomes were only detected in RNA isolated from BECs, and not in RNA isolated from neutrophils or Th17 cells. The expression of the BEC interferon response module correlated with viral genome copy number across all co-culture conditions indicating that the synergistic increased gene expression may be due to increased viral replication in BECs primed with Th17 cells. (**Figure 2E**)

*Th17 lymphocytes increase expression of IFN-signaling and viral response modules and decrease expression of cytoskeleton, intracellular transport, and RNA modification and splicing modules when co-cultured with RV-16 infected BECs*.

In parallel with BEC gene expression, RNA-sequencing was performed on RNA from Th17 lymphocytes after 5 days of co-culture with uninfected BECs or Th17 lymphocytes after 5-days of co-culture with RV-infected BECs (infected with RV-16 on day 4 of co-culture). (**Figure 1**)

Differential gene expression analysis identified 1899 genes differentially expressed in Th17 lymphocytes after infection of co-cultured BECs. Modular analysis identified 2 modules with decreased expression and 2 modules with increased expression after RV-16 infection of co-cultured BECs. (**Figure 3A**) The module with the greatest increase in expression was enriched for genes involved in response to virus and interferon signaling which we termed “Response to Virus and Interferons” that included the genes *STAT1, IRF1, IFI16, STAT2, IFIH1, IRF7, IFIT1, OAS1/2, ISG20, IFITM1, IFI27, IFITM2, IFITM3, SP100*, and *TBK1*. (**Figure 3A**) Modules thatdecreased expression in Th17 cells after rhinovirus infection of co-cultured BECs included “Ion Transport”, “Positive Regulation of Cellular Activation” and “Protein Localization.” (**Figure 3A**)

**Figure 3.**
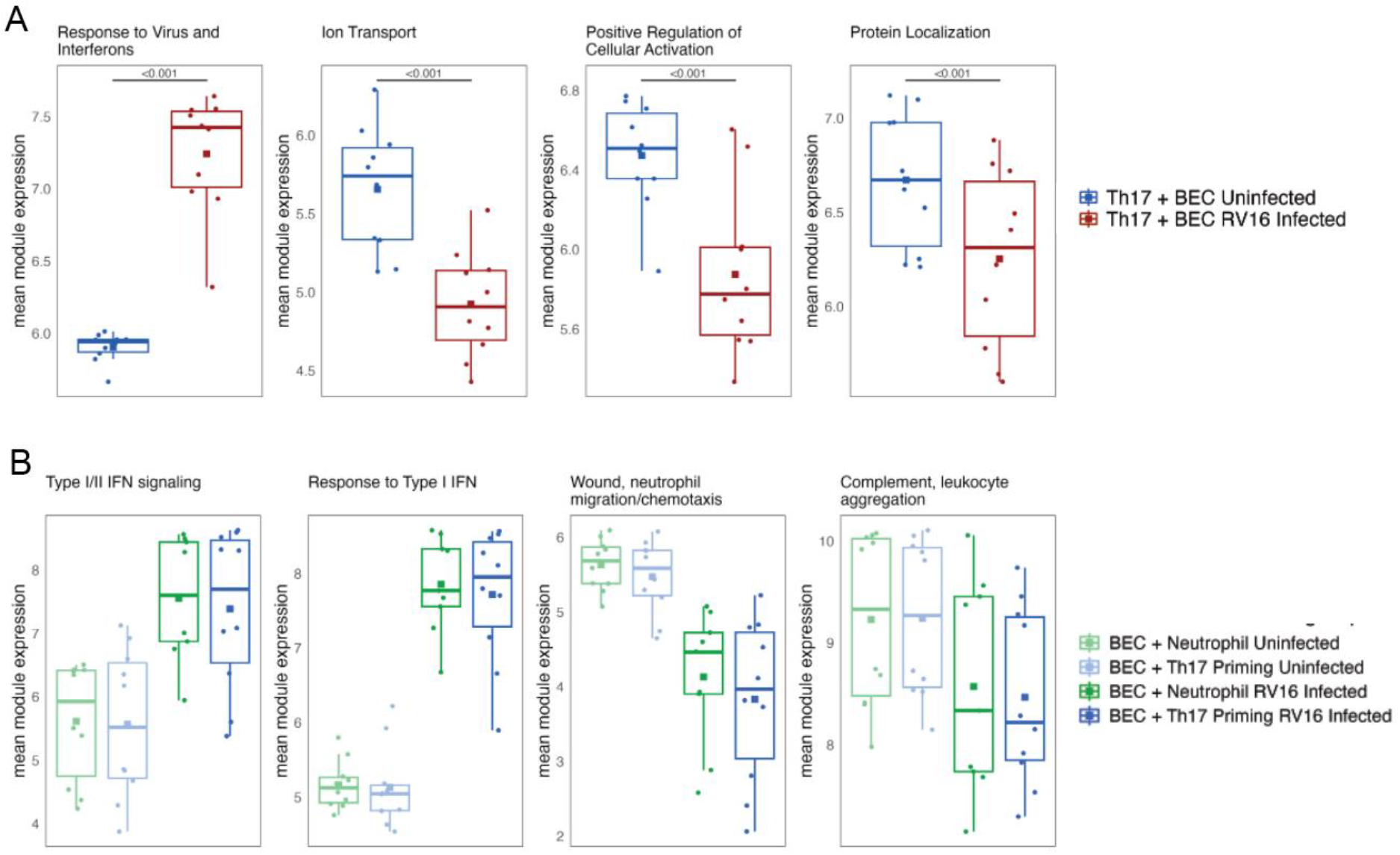
Infection of BECs with RV-16 induces transcriptional responses in co-cultured Th17 and neutrophils. **A:** Mean gene expression of 4 identifiable modules of gene expression in Th17 lymphocytes co-cultured with uninfected BECs for 5 days or for 5 days with BECs infected on day 4 of co-culture. **B.** Mean gene expression of 4 modules identified in neutrophils co-cultured with BECs for 24 hours either with naive BECs or Th17 primed BECs with and without RV-16 infection. All modules showed statistically significant differential expression when comparing uninfected to RV-16 infected conditions. For figure legibility, only the statistical comparisons within uninfected or within RV-16 infected conditions are shown.

### Neutrophil transcriptional responses to BEC infection and Th17 priming

Neutrophils were included after Th17-BEC co-culture as a downstream effector cell for Th17-BEC crosstalk. RNA was collected from neutrophils co-cultured for 24 hours with uninfected BECs or infected BECs that had previously been primed with 5 days of Th17 co-culture compared to primary BECs without priming. (**Figure 1**) Differential gene expression analysis identified 258 genes differentially expressed in neutrophils following viral infection, and no genes differentially expressed due to Th17 priming of BECs. (**Figure 3B**) Modular analysis identified 4 distinct gene modules with two exhibiting increasing gene expression and two exhibiting decreasing gene expression with RV infection of BECs. The two modules with increasing expression were composed from 91 and 52 genes respectively, with the first module named “Type I/II Interferon Signaling” including genes such as *IFITM1, BST2, IFITM3, IFI16, OAS2, SP100, IFIH1, IFI35, IRF7*, and *ISG20* and the second module named “Response to Type I Interferon” containing genes such as *TRIM5, MX1, MX2, IRF9, STAT1, TRIM56, TRIM5, ISG15, IFIT2, IFIT3, IFI27, IFITM2, IFI44L, OAS1, OAS3, IFI44*, and *HERC5*.

### Expression of interferon/viral response modules by Th17 lymphocytes correlates with expression of interferon signaling modules by BECs after RV infection of BECs

Analysis of module expression by BECs and Th17 lymphocytes showed significant positive correlation following RV infection of BECs, particularly for response modules in Th17 cells. (**Figure 4A&B**) Expression of the BEC module for Interferon Signaling and the Th17 module for Response to Virus and Interferons did not correlate in the uninfected state, but following RV infection expression increased in both cell populations and expression was highly correlated across each co-culture pair. (**Figure 4C**)

**Figure 4.**
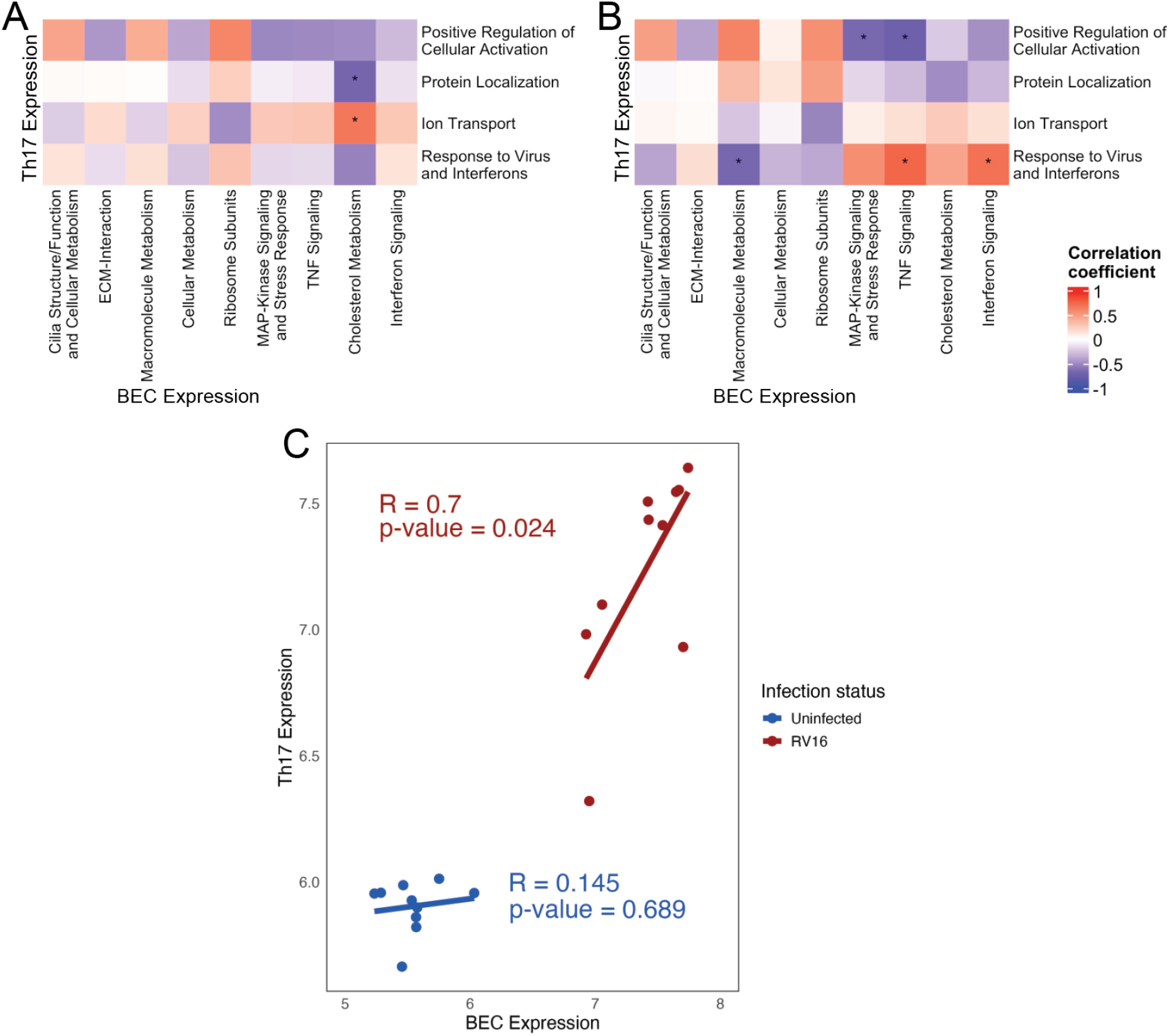
RV-16 infection leads to correlated gene expression in BECs and co-cultured Th17 cells. **A.** Correlation of module expression in co-cultured Th17 lymphocytes and uninfected BECs. ⋆p-value<0.05. **B**. Correlation of module expression in co-cultured Th17 lymphocytes and RV-16 infected BECs. ⋆p-value<0.05. **C**. Correlation of Interferon Signaling module in BECs with Interferon and Viral Response Module in Th17 lymphocytes with (red) and without RV-16 infection (blue).

### Th17 priming of BECs alters cytokine secretion following RV-16 infection of BECs

Measurement of secreted cytokines showed an altered secretion of IL-1β, CCL4, and CCL28 from Th17-primed BECs following RV-16 infection with IL-1β and CCL4 showing a synergistic increase following RV-16 infection of Th17 primed BECs. (**Figure 5A and D**) TNF-α and IL-17C increased with RV-16 infection. (**Figure 5C and E**) As BECs, Th17, and neutrophils all had a module of gene expression increasing after RV-16 infection consistent with an interferon-response set of genes, we measured protein levels of IFN-α and IFN-γ which both increased following RV-16 infection with an additive increase due to co-culture and RV-16 infection. (**Figure 5F and G**) Protein levels of CCL4, IL-1β, and TNF-α correlated with expression of the interferon signaling module in BECs with increased protein levels after RV-16 infection correlating with increased expression post-infection. (**Figure 6A-C**) TNF-α protein levels also correlated with increased expression of a MAP-kinase signaling module in BECs with Th17 co-culture as compared to BECs alone. (**Figure 6D**)

**Figure 5.**
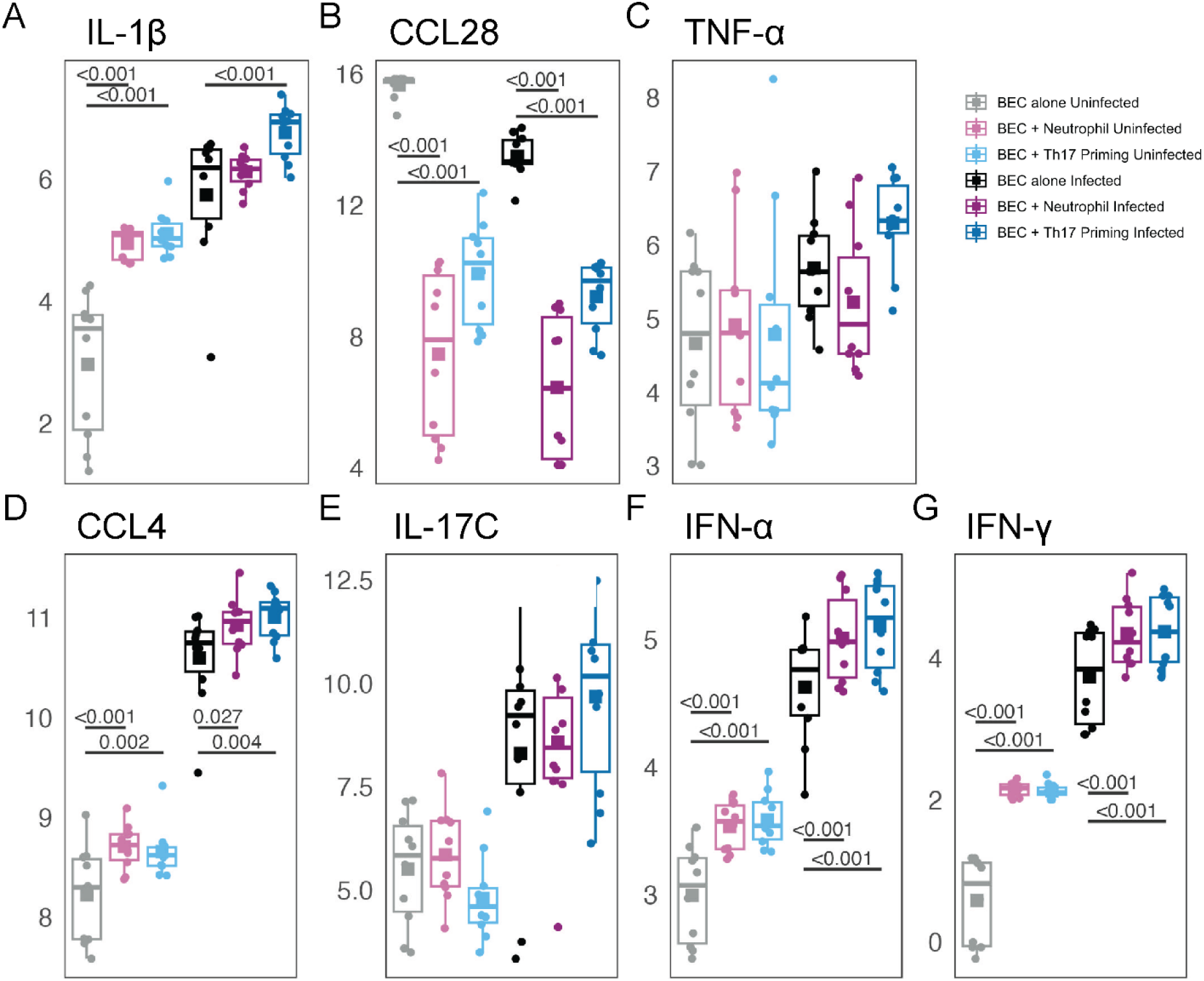
Th17 priming of BECs alters cytokine secretion following RV-16 infection of BECs. Secreted protein measurement of cytokines from each culture and infection condition were measured by Luminex protein quantification for IL-1β **(A)**, CCL28 **(B)**, TNF-α **(C)**, CCL4 **(D)**, IL-17C **(E)**, IFN-α **(F)**, and IFN-γ **(G)** measured as pg/mL. All proteins showed statistically significant differential expression when comparing uninfected to RV-16 infected conditions. For figure legibility, only the statistical comparisons within uninfected or within RV-16 infected conditions are shown.

**Figure 6.**
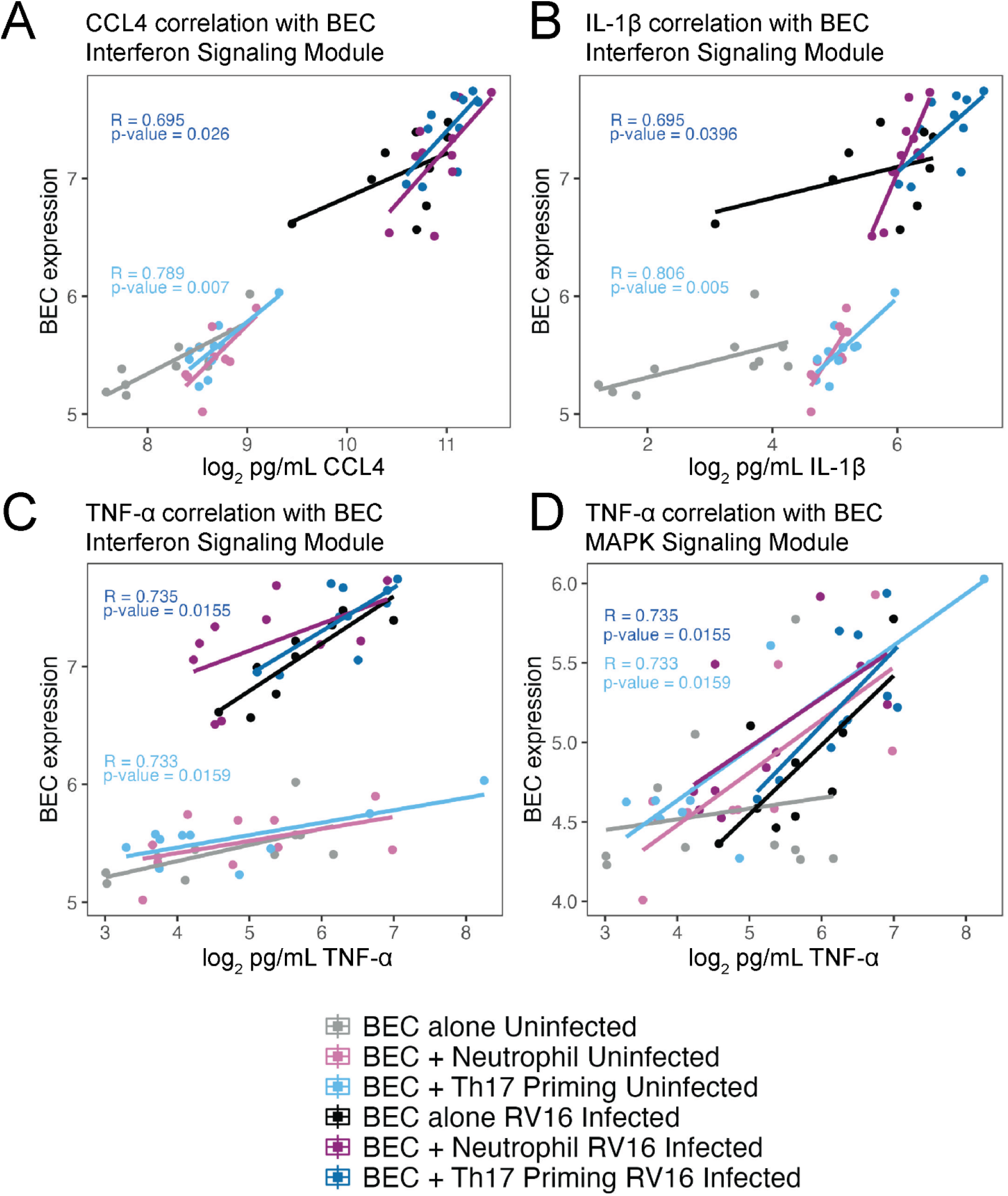
Gene expression in BECs correlates with secreted protein levels. **A.** Correlation of CCL4 protein levels in supernatant with BEC Interferon signaling module. **B**. Correlation of IL-1β protein levels in supernatant with BEC Interferon signaling module. **C**. Correlation of TNF-α protein levels in supernatant with BEC Interferon signaling module. **D**. Correlation of TNF-α protein levels in supernatant with BEC MAPK signaling module.

## Discussion

We investigated how Th17 co-culture alters gene expression in primary bronchial epithelial cultures from children with asthma and modifies the epithelial response to viral infections. We demonstrate that Th17 priming altered expression of six gene modules in RV-infected BECs. An interferon signaling module was synergistically activated by Th17 priming before RV infection and was associated with greater epithelial viral load. Th17 cells upregulated interferon-response genes when co-cultured with RV-infected epithelium, and BEC interferon signaling correlated with both Th17 interferon responses and viral copy number. Enhanced interferon signaling, together with elevated CCL4 and IL-1β secretion, indicated an enhanced innate epithelial response to RV-16 after Th17 priming. Our study provides several novel insights into epithelial–immune cell interactions relevant to non-T2 asthma and viral-triggered exacerbations.

We show that Th17 cells can reshape epithelial responses to RV infection leading to increased viral load and amplified post-infection antiviral interferon signaling in the epithelium. The differential gene expression we observed by RV-infected BECs with Th17 co-culture was likely driven by paracrine or innate immune signaling from Th17 cells as the priming occurred without specific activation of the Th17 cells. The Th17 cells were non-autologous cells, were never in direct contact with BEC, and were not directly stimulated or activated. Nevertheless, we observed many genes with altered expression in BECs under co-culture conditions alone and following RV infection. We observed that RV-16 genome copy number was significantly greater at 48 hours in RV-infected BECs primed by 4 days of Th17 co-culture as compared to RV-infected BECs cultured alone or with neutrophils. The increased viral copy number correlated with and mediated a synergistic increased expression of an interferon response module by epithelium conditioned by Th17 co-culture. Our observations contrast with earlier work using immortalized epithelial cell lines, where IL-17A signaling was reported to suppress rhinovirus replication.^21^ The contrasting observations may be due to inherent differences between transformed cell lines and primary organotypic epithelial cultures from asthmatic children, which more closely model *in vivo* biology and multiple signals from Th17 cells rather than the isolated impact of IL-17A alone.

Our findings using primary BEC cultures from children with asthma indicate that Th17 conditioning enhances epithelial viral replication leading to increased interferon-response gene expression in both BECs and Th17 cells, likely through a feedforward crosstalk mediated by secretion of IFN-α, IL-1β, and TNF-α from BECs. Such an interaction leading to increased viral replication and increased secondary interferon signaling could partially explain the risk for RV-triggered exacerbation in children with Th17-mediated asthma. In addition to the synergistic increase in interferon response genes, we observed an additive enhancement in expression of a MAP-kinase signaling and stress response module and a TNF-signaling module. Included in the MAP-kinase module was IL15 which has been inversely correlated with lung function in patients with severe asthma, and our finding that Th17 cells increased expression in the epithelium may provide a possible mechanism.^22^ The additive responses likely reflect the enhancement of an epithelial response in the setting of Th17 priming and highlight how epithelial-immune cell interplay can amplify an initial stimulus in the epithelium. The reciprocal correlation between interferon module expression in BECs and Th17 cells suggests a bidirectional signaling axis that may escalate inflammation during viral infections as Th17 priming increased epithelial expression of IL17RA which may poise the epithelium to respond to Th17 derived IL17 signaling. The increased interferon and inflammatory module expression in Th17 cells and BECs also correlated with increased measured levels of IFN-α, CCL4, and IL-1β following infection. Our demonstration that Th17 co-culture enhanced expression of TNF and MAP-kinase signaling modules further underscores the inflammatory amplification role of Th17–epithelial interactions. These pathways are strongly implicated in epithelial injury and remodeling, which are key features of asthma pathogenesis.^23–27^

In addition to an increase in inflammatory signaling, we found that BECs simultaneously downregulated cellular metabolism and cilia-associated gene modules with Th17 priming. Such a decrease in expression may impair epithelial barrier function and mucociliary clearance, thereby creating a permissive environment for viral propagation and contributing to mucus plugging in airways. Such changes could mechanistically explain clinical observations linking non-T2 asthma and neutrophilic inflammation with poor viral clearance and severe exacerbations. Importantly, our work provides evidence that Th17 cells dynamically respond to signals from epithelium infected with virus. Th17 cells have primarily been linked to bacterial and fungal responses, but here we find an interferon and viral response signal when co-cultured with RV-infected epithelium, despite the absence of detectable viral genomes within Th17 cells themselves. This observation indicates that soluble mediators, rather than direct infection, drive Th17 antiviral transcriptional programs. We demonstrate that Th17 cells can adopt a strong antiviral interferon related gene expression signature in response to epithelial infection, positioning them as amplifiers rather than bystanders in rhinovirus-triggered asthma exacerbations. As Th17 cells can have a quiescent or activated state, the increased expression of genes such as STAT1 or IRF7 in Th17 cells may indicate a transition to an activated inflammatory Th17 cell state, but this would need to be further investigated in future studies.

Inclusion of primary neutrophils in our BEC-immune cell co-culture experiments further extend insights from this work. Although Th17 priming of BECs did not directly alter neutrophil gene expression profiles, RV infection of BECs triggered strong interferon and viral response modules in neutrophils, similar to the responses observed in Th17 cells. One limitation of our design is that the Th17 and neutrophils were not in co-culture together, so we are unable to evaluate the impact of Th17 cells on neutrophils directly.

We acknowledge several other limitations in this study. The modest sample size (10 pediatric asthma donors) and use of non-autologous adult Th17 cells and neutrophils limit generalizability, and it remains unknown whether Th17 cells from asthmatic children would show greater dysregulation. The static 48-hour timepoint also precludes assessment of dynamic changes. Future work using sequential sampling before and after viral infection could also elucidate the kinetics of the responses found here as our analysis is limited to measurement of gene expression at 48 hours after infection. A further limitation is the absence of a Th17-only priming condition, which would have clarified BEC-specific effects but is less representative of in vivo airways where Th17 and IL-17A signaling contribute to baseline neutrophilic inflammation. Despite these limitations, the use of primary, organotypic epithelial cultures from well-characterized children with asthma represents a major strength, providing physiologically relevant insights that extend beyond transformed cell lines. Finally, the sample size prevented correlation with donor clinical measures such as asthma severity, prior exacerbations, or T2-high/non-T2 phenotype.

A novel aspect of our study is the demonstration that Th17 priming potentiates epithelial secretion of proinflammatory cytokines following RV infection. Secreted IL-1β, CCL4, IFN-α and IFN-γ were increased by Th17 priming of BECs prior to infection. These cytokines not only serve as markers of antiviral and inflammatory activity but also act as upstream regulators of broader immune cascades. Elevated IL-1β and CCL4 can promote epithelial damage, recruit neutrophils, and perpetuate inflammation, while increased IFN-α may further activate interferon-stimulated gene networks in both epithelial and immune cells. Thus, our findings provide mechanistic insight into how Th17 activity may set the stage for self-reinforcing inflammatory responses during viral exacerbations of asthma. Together, these findings advance our understanding of inflammatory responses in RV-triggered exacerbations in non-T2 asthma in several important ways. First, they support the hypothesis that Th17-mediated infection pathways, rather than being solely antibacterial or antifungal, play a central role in shaping epithelial responses to viral pathogens. Second, they suggest that the presence of Th17 cells in the asthmatic airway fundamentally alters the outcome of viral infection, leading to increased viral load and heightened inflammation. Third, they highlight the importance of epithelial–Th17 crosstalk as a driver of neutrophilic inflammation and impaired epithelial homeostasis, processes that are hallmarks of severe, corticosteroid-refractory asthma. These mechanistic insights may help explain why children with Th17-associated asthma phenotypes experience frequent, severe viral exacerbations despite standard therapy targeting T2-inflammation.

Our study also identifies potential important therapeutic implications. Targeting IL-17 signaling has been explored in asthma, but clinical trials to date have shown limited benefit, possibly because blocking IL-17 alone does not fully address the complex epithelial–Th17– neutrophil network. Our data suggest that interventions designed to modulate epithelial responses to Th17 signals, limit viral replication, or dampen synergistic cytokine amplification (e.g., TNF-α or IL-1β blockade) may hold promise for reducing exacerbation severity in non-T2 asthma. Furthermore, enhancing epithelial resilience, for example through therapies that restore ciliary function or metabolic integrity, may counteract the permissive effects of Th17 priming on viral replication.

In conclusion, our study reveals that Th17 lymphocytes profoundly shape epithelial and immune responses to rhinovirus infection in the pediatric asthmatic airway. Th17 priming enhances viral replication, amplifies interferon and proinflammatory gene programs in both epithelium and Th17 cells. This bidirectional cross-talk establishes a feed-forward loop of viral propagation and inflammation that may underlie the heightened susceptibility to viral exacerbations in both T2 and non-T2 asthma. By identifying epithelial–Th17 interactions as critical regulators of viral responses, our findings advance mechanistic understanding of non-T2 asthma and highlight novel pathways that may be targeted to prevent or attenuate severe viral-induced exacerbations in children, particularly in children who have not responded to therapies targeting T2-inflammation.

## Supplemental Figures and Tables

**Supplemental Table E1**. List GenBank accession numbers for custom viral reference database.

**Supplemental Table E2**. Full model results of BEC gene expression by comparisons for co-culture and infection status.

**Supplemental Table E3**. Module composition for BECs.

**Supplemental Table E4**. Module composition for Th17 cells

**Supplemental Table E5**. Module composition for neutrophils.

**Supplemental Table E6**. Protein levels by co-culture and infection status.

**Supplemental Table E7**. Correlation of protein levels with BEC module expression.

## Supplemental Figures

**Figure E1.**
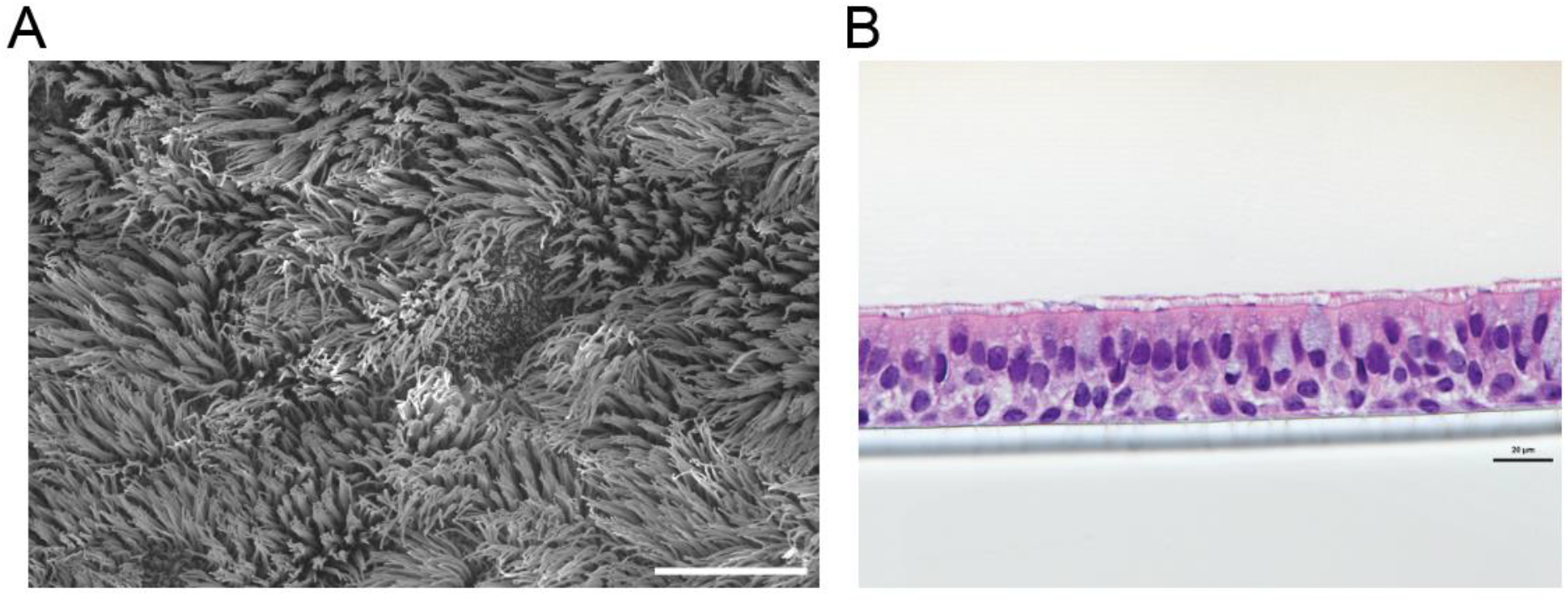
**A**. Scanning electron microscopy of a representative mature BEC culture showing ciliated apical layer. Scale bar 10 microns. **B**. H&E staining of a representative mature BEC culture with pseudostratified epithelium. Scale bar 25 microns.

